# Persistence and anti-persistence in treadmill walking

**DOI:** 10.1101/2021.04.12.439523

**Authors:** Klaudia Kozlowska, Miroslaw Latka, Bruce J. West

## Abstract

**Background:** Long-range persistent correlations in stride time (ST) and length (SL) are the fundamental traits of treadmill gait. Our recent work showed that the ST and SL time series’ statistical properties originated from the superposition of large-scale trends and small-scale fluctuations (residuals). Trends served as the control manifolds about which ST and SL fluctuated. The scaling exponents of the residuals were slightly smaller than 0.5.

**Research question:** Do random changes in treadmill belt speed affect the trend properties and scaling exponents of ST/SL residuals?

**Methods:** We used Multivariate Adaptive Regression Splines (MARS) to determine gait trends during a walk on a treadmill whose belt speed was perturbed by a strong random noise. Then, we calculated the scaling exponents of MARS residuals with the madogram estimator.

**Results:** Except for the ST at the lowest treadmill speed *v* = 0.8 m/s, the normalized trend duration was at least three times greater than that for the unperturbed walk. The Cauchy distribution scale parameter, which served as a measure of the width of SL and ST trend slope distributions, was at *v* = 1.2 m/s, almost 50% and 25% smaller than the unperturbed values. The differences were even greater at *v* = 1.6 m/s: 73% and 83%. For all speeds, the ST and SL MARS residuals were strongly anti-persistent. At *v* = 1.2 m/s, the corresponding scaling exponents were equal to 0.37±0.10 and 0.25±0.09. Apart from ST at *v* = 0.8 m/s, the ST/SL scaling indices were close to 0.5.

**Significance:** Persistence of gait parameters is closely related to the properties of their trends. Longer trends with a gentle slope and strong anti-persistence of ST/SL residuals are the manifestations or tight control required during the perturbed treadmill walk.

## 1. Introduction

Motor control is the process of initiating, directing, and grading a purposeful voluntary movement. It is surprising how many studies on variability in movement execution contributed to the understanding of motor control and skill learning [1, 2, 3]. Woodworth’s classic paper [4] on line drawing published in 1899 pioneered this consequential line of research. Fluctuations in gait spatio-temporal parameters such as stride time (ST), stride length (SL), and stride speed (SS) attracted general interest almost a century later due to the unexpected discovery of long-range, persistent correlations in human ST [5, 6]. Shortly after that, such correlations were observed in sequences of ECG RR intervals [7], inter-breath intervals [8], and beat-to-beat cerebral blood flow [9], to name a few. Physiological time series persistence was at variance with the homeostasis considered at that time the guiding principle of medicine [10]. On the other hand, the findings corroborated the emerging concept of fractal physiology [11], which shifted the emphasis from the mean values of parameters to their variability. Thus, not unexpectedly, Hausdorff’s et al. papers [5, 6] have spawned a flurry of research on the origins and significance of correlations in human gait in particular and in physiological time series in general.

Some argued that persistent fluctuations manifest the adaptability of underlying control systems and the absence of long-range correlations or anti-persistence indicates disease or pathology [12, 13, 14]. Deligniéres and Torre demonstrated that when healthy humans walk in sync with a metronome, their stride times become anti-persistent [15]. During treadmill walking, a subject’s SS must fluctuate about the treadmill belt’s speed. It turns out that, unlike ST and SL, SS is anti-persistent. Moreover, all three parameters are anti-persistent during treadmill walking with auditory [16] or visual [17] cueing (alignment of step lengths with markings on the belt). Thus, anti-persistence can not only be a manifestation of pathology but also of tight control [18, 19, 20, 21].

Our recent work [22] shows that during treadmill walking, the statistical properties of ST and SL time series originate from the superposition of large-scale trends and small-scale fluctuations (residuals). Trends serve as the control manifolds about which SL and ST fluctuate. The trend speed, defined as the ratio of instantaneous values of SL and ST trends, is tightly controlled about the treadmill speed. The trends’ strong correlation ensures that their concomitant changes result in a movement along the constant speed goal equivalent manifold [23, 20]. The scaling exponents of ST and SL residuals are slightly smaller than 0.5, indicating weak anti-persistence.

For healthy subjects, walking on a treadmill is straightforward. The question arises as to whether the properties of trends and residuals change when speed control becomes more challenging. For example, when the belt speed randomly varies. We hypothesize that the tighter control of SL and ST, which is required to compensate for perturbations, results in smaller trends’ slopes, longer trend’s duration, and strong anti-persistence of the residuals.

## 2. Methods

### 2.1. Experimental Data

In our analysis, we used a dataset from the study of Moore et al. [24] that is available from the Zenodo repository [25]. Fifteen young, healthy adults walked on a motor-driven treadmill for ten minutes at three speeds (0.8 m/s, 1.2 m/s, and 1.6 m/s). Each trial started with a one-minute normal (unperturbed) walk (first normal walk – FNW). This phase was followed by an 8-minute longitudinally perturbed walk (PW). The trial ended with a 1-minute second normal walk (SNW). Due to technical issues, we discarded some records. The analyzed dataset consisted of 11, 12, and 10 records for treadmill speed 0.8 m/s, 1.2 m/s, and 1.6 m/s, respectively. In the PW part of the trial, during each stance phase of the subject’s gait, the treadmill belt’s speed varied randomly. The standard deviation of the noise was equal to 0.06 m/s, 0.12 m/s, and 0.21 m/s for 0.8 m/s, 1.2 m/s, and 1.6 m/s experiments, respectively. A comprehensive description of the experimental protocol and participants’ characteristics can be found in the original paper.

The motion capture trajectories of RLM and LLM (right/left lateral malleolus of the ankle) markers were resampled at 100 Hz to determine a subject’s stride length (SL), stride time (ST), and stride speed (SS). A heel strike was defined as the point where the forward foot marker was at its most forward point during each gait cycle. Step duration was equal to the elapsed time between the ipsilateral and contralateral heel strikes. We calculated SL and ST as the sum of the corresponding values for two consecutive steps. The quotient of SL and ST yielded SS.

### 2.2. Identification of trends in gait parameters

We used Multivariate Adaptive Regression Splines (MARS) [26], to approximate trends in gait spatio-temporal parameters. We employed a piecewise linear, additive MARS model (the order of interaction was equal to 1) with a maximum number of basis functions equal to 50. The generalized cross-validation knot penalty *c* and the forward phase’s stopping condition were set to 2 and 0.001, respectively. A detailed description of MARS gait trend analysis can be found in our previous paper [27].

A normalized trend duration, *TD*, is defined as the trend’s duration divided by the subject’s mean stride time. A normalized trend slope *TS* is the change of the gait parameter divided by the product of its average value and the normalized trend duration.

The Cauchy distribution’s scale parameter *γ* serves as the measure of the width of SL and ST trend slope distributions [27]. Please note that the expected value and variance of the Cauchy distribution are undefined [28].

We analyzed the statistical properties of trends with Mathematica 12.1 (Wolfram Research).

### 2.3. Scaling analysis

In their seminal papers, Hausdorff et al. [5, 6] applied detrended fluctuation analysis (DFA)[29] to quantify correlations in human stride time. The choice of DFA implied that a stride duration time series was made up of persistent (scaling exponent greater than 0.5) fractal fluctuations superposed on trends that are irrelevant from the point of view of fractal analysis. Over the last quarter-century, this was the method of choice for determining the gait scaling exponents. In DFA, the signal is divided into analyzing windows of different sizes in which a trend (typically a polynomial of a priori chosen order) is removed. Partitioning the signal into a set of nonoverlapping windows and performing detrending in a window-based manner does not guarantee that the functional form of the trend in each window is identical to the assumed one. This is especially true for small windows. Contrary to expectations, this critical finite-size effect is always present [30, 31]. The only solution is to apply adaptive detrending such as MARS. In our recent work, we demonstrated that DFA is incapable of removing gait trends [27]. Phinyomark et al. [32] warn against using DFA for the interpretation of scaling properties of gait time series.

In this work, we employ the madogram estimator (MD) to calculate the gait scaling exponents.

The variogram of order *p* of a stochastic process with stationary increments is defined as one half times the expectation value of an increment at lag *t* [33]:

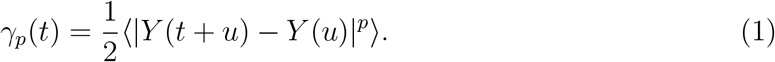

For *p* = 1 we call such a structure function the madogram. As *t* → 0

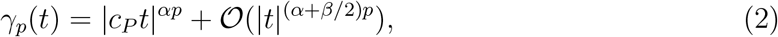

where *H* ∈ (0, 1] and constants *β* and *c*_*P*_ are positive. For one-dimensional time series 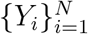, we may define the power variation of order *p*:

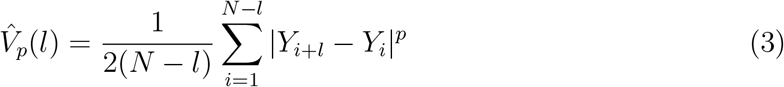

and using Eq. (2) derive the following estimate of fractal dimension:

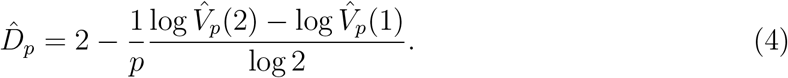

The fractal dimension 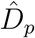 and scaling exponent *α* are related by the following equation:

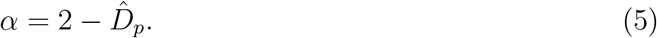

The madogram estimator (*p* = 1) turns out to be particularly robust, especially for non-Gaussian processes. Gneiting er al. [33] compared the properties of fractal dimension estimators that were implemented in the R package fractaldim. The following MATLAB function implements the madogram:

~~~
function [alpha] = madogram(x)
% calculates the Hurst exponent (alpha) of vector x using the madogram
% Statist. Sci. 27(2) : 247-277 DOI : 10.1214/11-STS370
% Use MATLAB’s wfbm function for testing : madogram (wfbm (0.7, 2048))
   n=length(x);
   V1=0;
   V2=0;
   for i=1:n-1
         V1=V1+(abs(x(i+1)-x(i)));
   end
   for i=1:n-2
         V2=V2+(abs(x(i+2)-x(i)));
   end
   V1=0.5∗V1/(n-1);
   V2=0.5∗V2/(n-2);
   alpha=(log (V2)-log (V1))/log(2);
end
~~~

We performed the scaling analysis for the raw experimental time series and time series detrended using the piecewise linear variant of the MARS model. In the latter case, the exponents have an *L* subscript (*α*^(*L*)^).

### 2.4. Trend speed

A trend speed 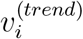 is the ratio of the values of piecewise linear SL and ST MARS trends at i*th* stride. To quantify the deviation of the trend speed from the average stride speed *< SS >* during a given trial, we define a trend speed control (*TSC*) parameter as:

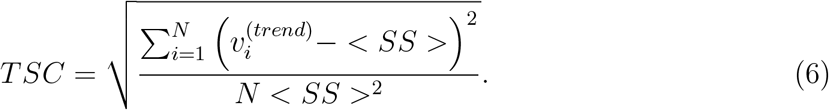

Thus, *TSC* is equal to zero when the trend and mean stride speed are equal during the whole trial.

### 2.5. Statistical analysis

We used the Shapiro-Wilk test to determine whether the analyzed data were normally distributed. The dependence of gait scaling indices on treadmill speed was examined with either ANOVA or the Kruskal-Wallis test (with Tukey’s post hoc comparison in both cases). For a given gait parameter and treadmill speed, the difference between *α* and *α*^(*L*)^ was assessed with either the *t*-test or the Wilcoxon signed rank test. The same tests were used to check whether the scaling indices were smaller than 0.5.

For all statistical tests, we set the significance threshold to 0.05.

In the presented boxplots, the black and grey dots correspond to the outliers and far outliers, respectively.

## 3. Results

Fig. 1 shows the examples of stride time, length and speed time series. The solid thick lines in this figure represent trends approximated using the piecewise linear variant of the MARS model.

**Figure 1:**
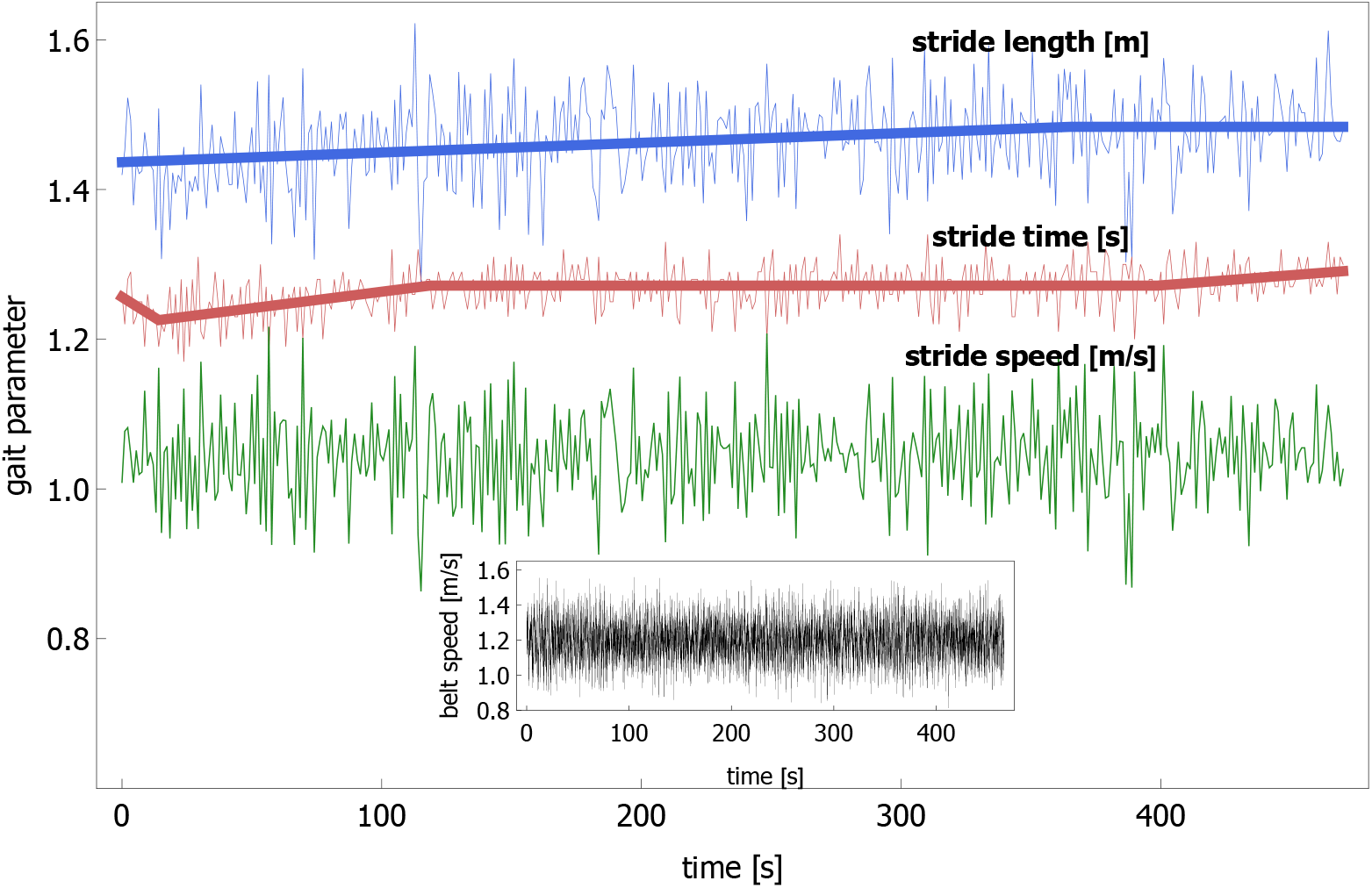
Stride length, time, and speed during the perturbed walk of subject 10 at 1.2 m/s. The data come from the study of Moore et al. [24]. For clarity, the stride time and length time series were vertically shifted by 0.15 s and 0.30 m, respectively. Thick solid lines show trends in stride time and length calculated using the piecewise linear variant of the MARS model. There were no trends in stride speed. The inset shows the instantaneous belt speed during the perturbation phase of the trial.

### 3.1. SL and ST trends

There were 129 SL trends (34, 45, and 41 for treadmill speed *v* equal to 0.8 m/s, 1.2 m/s, and 1.6 m/s, respectively) and 217 ST trends (110, 71, and 36). The SL and ST trends were strongly correlated and independent from the corresponding residuals. The Pearson correlation coefficient was equal to 0.73±0.12, 0.85±0.13, and 0.88±0.08 for the successive speeds, respectively. The difference in the correlation strength between the smallest and highest *v* was statistically significant (*p* = 0.04).

The trend duration *TD*_*SL*_ was speed independent (*p* = 0.67): 90±92, 100±122, and 123±131 for the successive *v*, respectively. In stark contrast, *TD*_*ST*_ increased with *v*±±(*p* = 1× 10^*-*5^): 38±41, 77±90, and 140±127 (these values were statistically different from each other).

For SL, the Cauchy distribution scale parameter *γ*_*SL*_ was equal to 7 × 10^*-*4^, 7 × 10^*-*4^, and 4 × 10^*-*4^ for 0.8 m/s, 1.2 m/s, and 1.6 m/s, respectively. For ST, the corresponding values of *γ*_*ST*_ were equal to 1.6 × 10^*-*3^, 9 × 10^*-*4^, and 2 × 10^*-*4^. While *γ*_*SL*_ did not depend on *v, γ*_*ST*_ at the smallest and highest *v* were statistically different.

Fig. 2 shows the best-fit Cauchy probability density function for ST and SL. We aggregated data from the experiments at the two highest treadmill speeds (1.2 m/s and 1.6 m/s). The probability plots, presented as insets, show the differences between the experimental and Cauchy’s cumulative distribution function (CDF). One can see that also in the presence of perturbations, the Cauchy distribution fairly approximates the experimental data.

**Figure 2:**
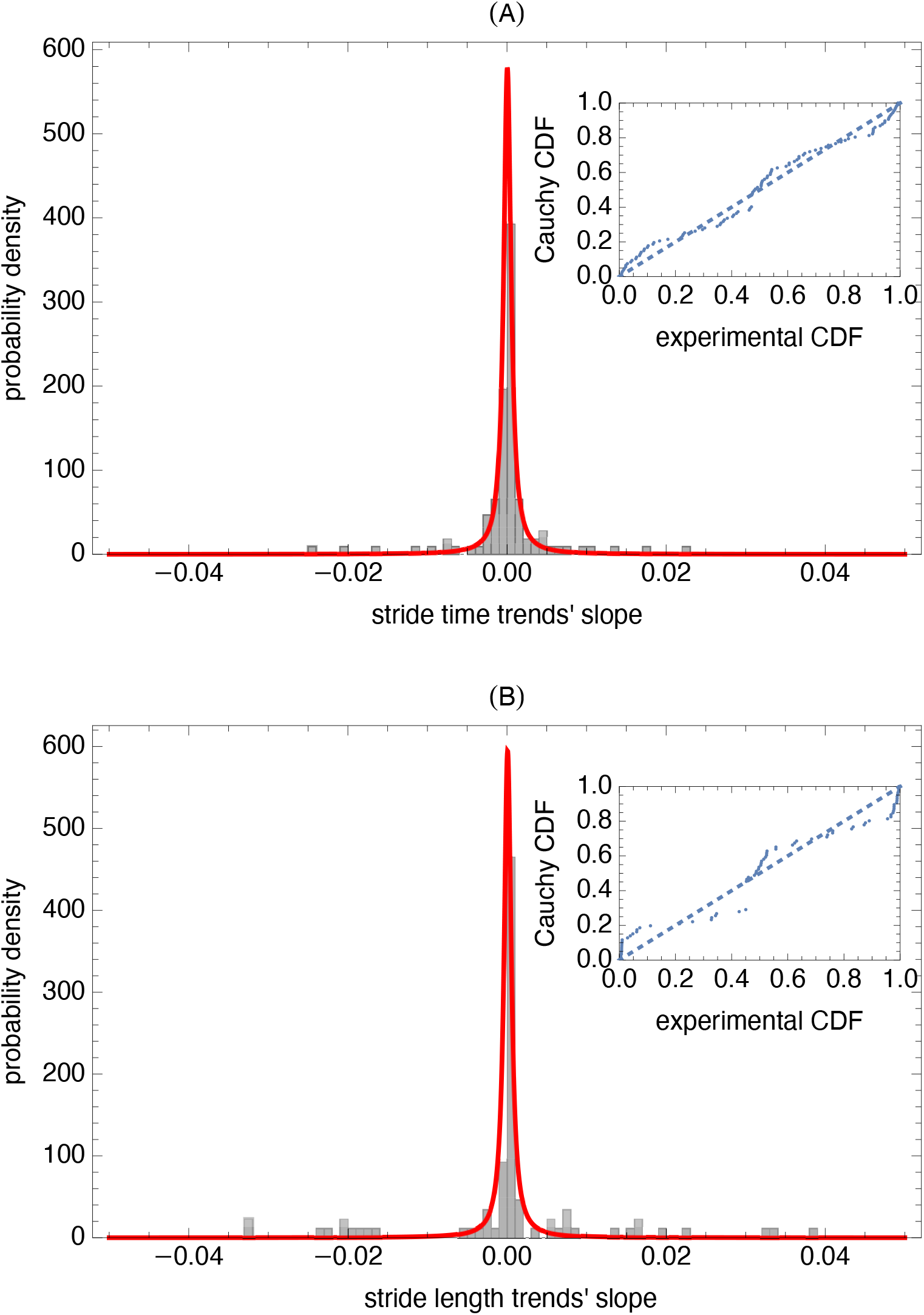
Probability density function (PDF) of normalized trend slopes of: (A) stride time and (B) stride length. We aggregated data from the experiments at the two highest treadmill speeds (1.2 m/s and 1.6 m/s). The solid curves in both subplots show the best-fit Cauchy PDFs. The probability plots, presented as insets, show the differences between the experimental and Cauchy’s cumulative distribution functions (CDFs).

### 3.2. Scaling exponents

Table 1 shows the mean value and standard deviation of ST, SL, and SS scaling exponents at different treadmill speeds. The boxplots in Fig. 3 visualize the distributions of *α* and *α*(^*L*^).

**Table 1:** Scaling exponents of spatio-temporal gait parameters for three treadmill speeds. The exponents were calculated for the experimental time series (*α*) and the MARS residuals (*α*^(*L*)^). Data are presented as mean±standard deviation. All indices that are not statistically smaller than 0.5 are marked with the dagger symbol.

| $\alpha_{SL}$ | $\alpha_{SL}^{(L)}$ | $\alpha_{ST}$ | $\alpha_{ST}^{(L)}$ | $\alpha_{SS}$ | $\alpha_{SS}^{(L)}$ |
| --- | --- | --- | --- | --- | --- |
| **speed 0.8 m/s** |  |  |  |  |  |
| $0.45 \pm 0.06$ | $0.39 \pm 0.09$ | $0.64 \pm 0.10^\dagger$ | $0.41 \pm 0.08$ | $0.28 \pm 0.10$ | $0.27 \pm 0.10$ |
| **speed 1.2 m/s** |  |  |  |  |  |
| $0.42 \pm 0.12$ | $0.37 \pm 0.10$ | $0.44 \pm 0.14^\dagger$ | $0.25 \pm 0.09$ | $0.15 \pm 0.10$ | $0.15 \pm 0.10$ |
| **speed 1.6 m/s** |  |  |  |  |  |
| $0.49 \pm 0.11^\dagger$ | $0.43 \pm 0.10$ | $0.42 \pm 0.09$ | $0.33 \pm 0.11$ | $0.28 \pm 0.08$ | $0.28 \pm 0.08$ |

**Figure 3:**
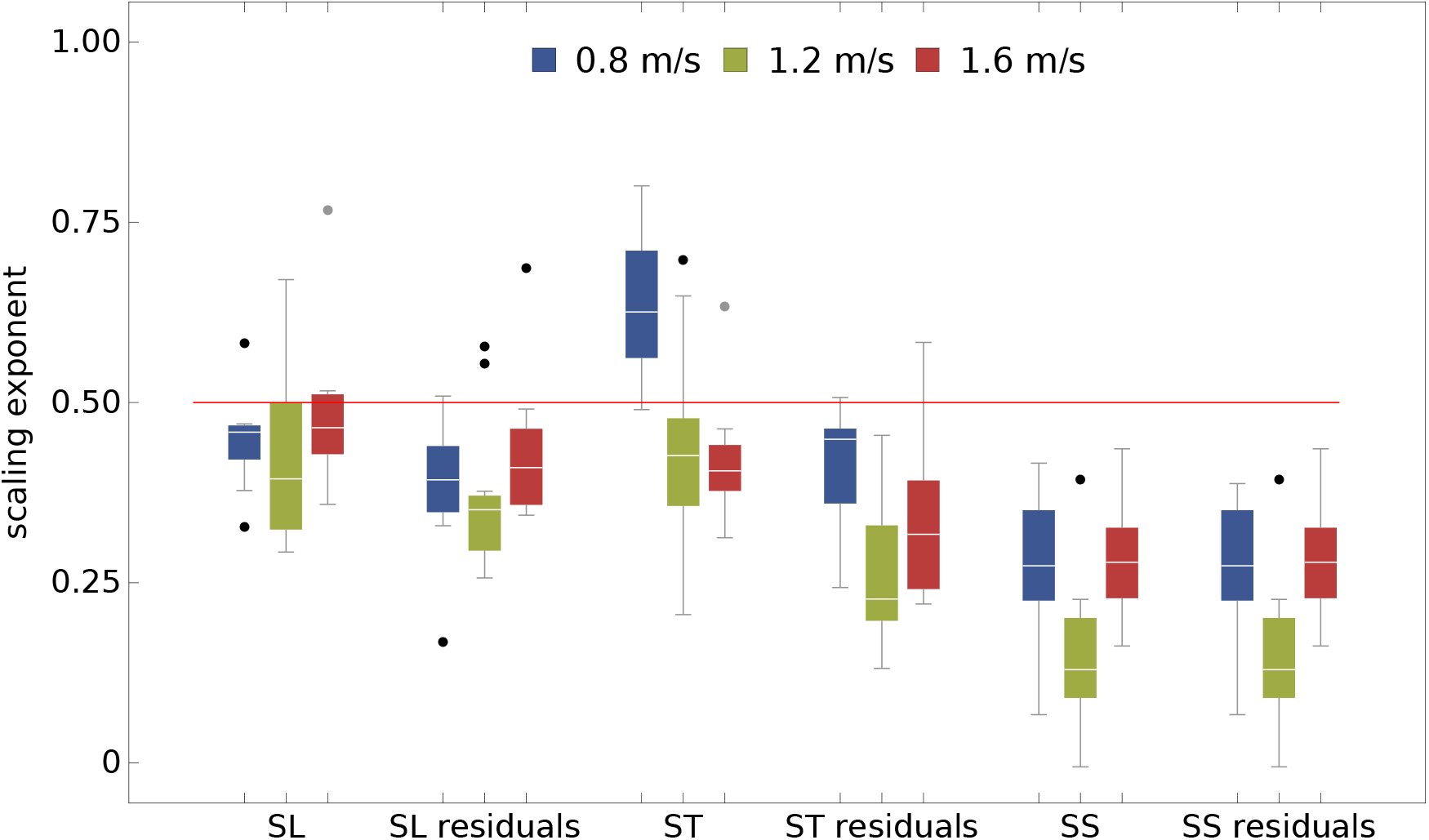
The boxplots of scaling exponent of spatio-temporal gait parameters for three treadmill speeds. The exponents were calculated for the raw experimental time series and the MARS residuals. The horizontal red line delineates persistent (*>* 0.5) and anti-persistent (*<* 0.5) scaling.

Except for *α*_*ST*_ at *v* = 0.8 m/s and *v* = 1.2 m/s as well as *α*_*SL*_ at *v* = 1.6 m/s, all exponents were statistically smaller than 0.5 – indicating anti-persistence of the experimental time series and their MARS residuals.

For SL and ST, regardless of treadmill speed, *α*^(*L*)^ was smaller than *α*. Detrending did not affect the SS scaling indices.

*α*_*SS*_ and 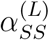 had a minimum at *v* = 1.2 m.s (value close to the preferred walking speed of adult subjects). Qualitatively, the same effect was observed for the SL and ST residuals. However, except for 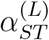 at the lowest treadmill speed, the differences between the value of the scaling exponent at *v* = 1.2 m/s and these for the smallest and highest velocity were not significant.

### 3.3. Trend speed

The boxplots of the trend speed control parameter *TSC* in Fig. 4 show that the control of *v*^*trend*^ increases with the treadmill speed: 0.04±0.03, 0.02±0.03, and 0.02±0.03, for the successive speeds, respectively. The difference in *TSC* between *v* = 0.8 m/s and *v* = 1.6 m/s is statistically significant (*p* = 0.005).

**Figure 4:**
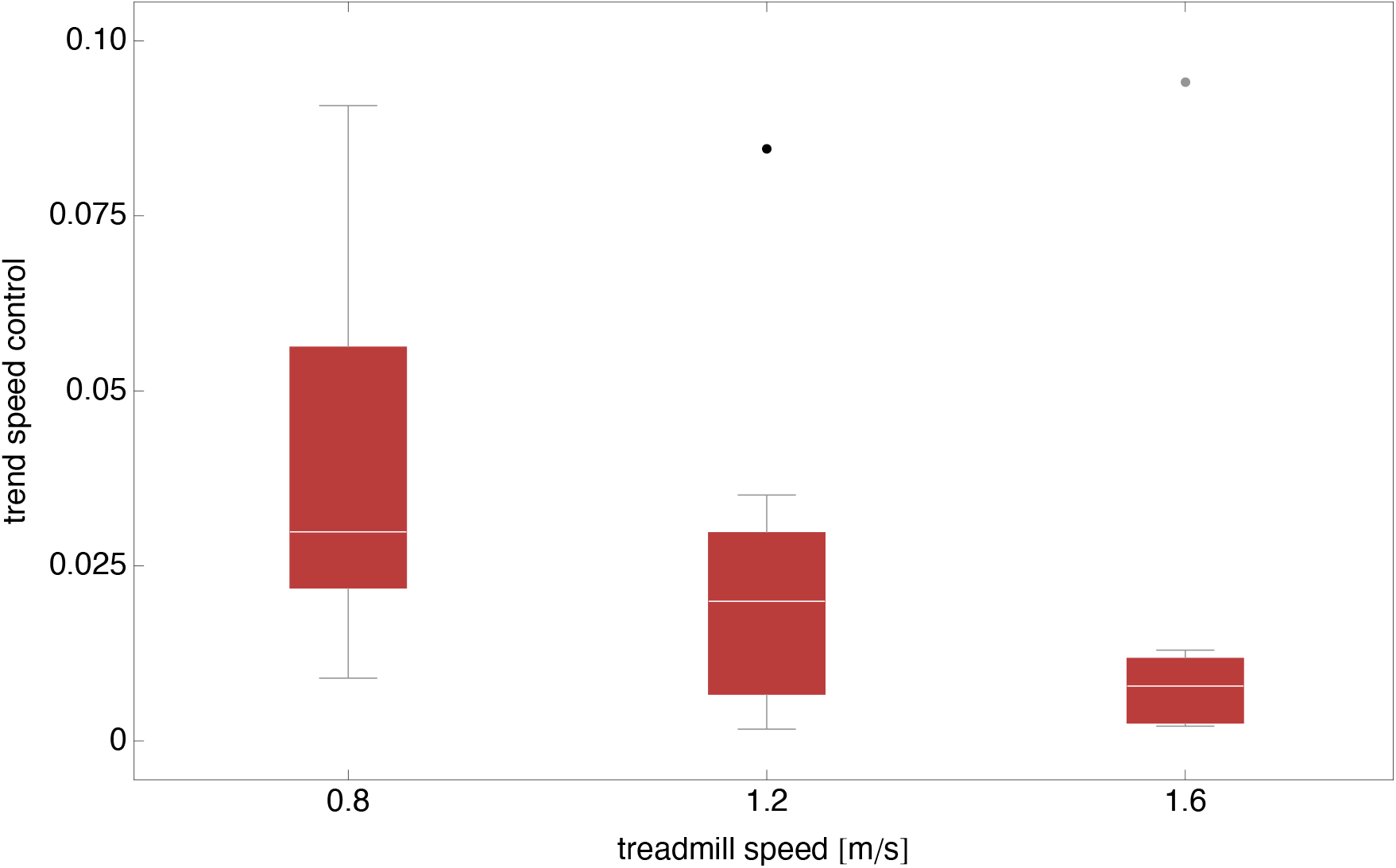
The boxplots of trend speed control parameter (*TSC*).

## 4. Discussion

Our previous study [22] showed that during an unperturbed walk in Dingwell et al. experiment [18], the distribution of trend duration and trend slope was speed independent and similar for ST and SL. We found that *TD*_*ST*_ = 23±26 and *TD*_*SL*_ = 25±36. The scale parameter *γ* of the Cauchy distribution fit to the trend slope distribution was equal to *γ*_*ST*_ = 1.2 × 10^*-*3^ and *γ*_*SL*_ = 1.5 × 10^*-*3^. Please note that we refer to Dingwell’s dataset because the unperturbed walking segments in Moore’s study [24] were too short for both trend and scaling analysis.

At the lowest treadmill speed *v* = 0.8 m/s, longitudinal perturbations have different effects on ST/SL trends. In particular, the number of ST trends was almost three times larger than that for SL (110 vs. 43). Moreover, the average *TD*_*ST*_ = 38±41 was much closer to the unperturbed values in comparison with SL: *TD*_*SL*_ = 90±92. It is plausible that at such a low *v*, only the SL adjustments suffice to compensate for the perturbations [34].

For higher *v*, the differences between ST and SL disappear. The trend duration and trend slope are drawn from the same distribution. As in unperturbed walking [27], the mean and standard deviation of *TD* are close to each other – the unmistakable sign of an exponential tail. It is worth emphasizing that except for the ST at the lowest speed, ST/SL *TD* is at least three times greater than the unperturbed value. The Cauchy distribution scale parameter *γ*, which we used as a measure of the width of trend slope distribution, at *v* = 1.2 m/s, was almost 50% (SL) and 25% (ST) smaller than the regular walk values. The differences were even greater at *v* = 1.6 m/s: 73% and 83%.

One could argue that the comparison between the unperturbed and perturbed walk is hindered by the fact that in Dingwell’s experiment, treadmill speed was varied as a percentage of their preferred walking speed (PWS). Therefore, we also reanalyzed data from our previous work [35] in which the subjects walked 400 m at three speeds: 1.1 m/s, 1.4 m/s, and 1.7 m/s. We will elsewhere show that the trend properties were very similar to those observed in Dingwell’s data.

Human voluntary movements are by their very nature redundant [2]. The sensorimotor system does not have to select a unique solution but rather samples the family of solutions congruent with the task. Regardless of skill level, no movement is ever the same; there is always variability. Bernstein recognized this property, which is also known as the dimensionality problem [36]. In his own words, there is ‘repetition without repetition.’ Todorov advocated that sensorimotor controllers obey a minimal intervention principle [37]. They act only on those variables in the multidimensional state space that are goal-relevant. The other variables are free to vary and form the so-called uncontrolled manifold (UCM) [38].

Dingwell et al. [19] realized that both the minimal intervention principle and energy optimization underlay the control of spatio-temporal gait parameters during treadmill walking. Generally, during such movement, any SL and ST whose ratio is equal to the belt’s speed lies on the UCM. However, humans minimize the energy cost of walking by choosing specific values of both parameters at a given speed. Thus, there is a preferred operating point (POP) on the constant speed goal equivalent manifold (GEM). Dingwell et al. used the SL and ST mean values as the POP components. They then projected the deviation vector onto the GEM and the axis perpendicular to it. It turns out that the tangential and transverse components are persistent and anti-persistent, respectively. In agreement with the optimal control theory, the tangential variability is higher than the transverse one. Thus, these statistical properties provide evidence that subjects do not regulate ST and SL independently but instead adjust them simultaneously to maintain a stable walking speed.

Herein and in our recent work [27], we demonstrated that trends serve as control manifolds about which ST and SL fluctuate. The strong coupling between the ST and SL trends ensures that their concomitant changes result in movement along the GEM. In the presence of perturbations, at *v* = 1.2 m/s, the trend speed control parameter *TSC* is about six times smaller compared to the regular treadmill walking (0.02 vs. 0.13, see Fig. 4) [27], reflecting the precise control of trend speed. During overground or regular treadmill walking, trends can be conspicuous. When tighter control is needed, for example, to compensate for random changes in treadmill belt speed, the trends become longer, flatter, and the ST/SL residuals more anti-persistent. For weak trends, even the scaling exponents of the experimental ST/SL time series *α*_*ST*_ and *α*_*SL*_ are small – the effect we observed in unperturbed walking [22]. One can see in Table 1, that except for *α*_*ST*_ at *v* = 0.8 m/s the ST/SL mean values are close to 0.5. In half of the cases, the experimental time series fluctuations are anti-persistent.

The presented findings corroborate our reinterpretation of the scaling properties of gait time series [22]. The persistence of stride time and length during treadmill walking is closely related to the properties of their trends. The strong ST-SL trend coupling is an automatic way of controlling gait speed and can be regarded as a manifestation of the minimum intervention character of human motor control.

